# An actomyosin-polarized membrane reservoir mediates the unequal divisions of *Drosophila* neural stem cells

**DOI:** 10.1101/2022.05.03.490551

**Authors:** Bryce LaFoya, Kenneth E. Prehoda

## Abstract

**Summary:** The asymmetric divisions of *Drosophila* neural stem cells (NSCs) produce unequally sized siblings, with most volume directed into the sibling that retains the NSC fate. Sibling size asymmetry results from preferential expansion of the NSC sibling surface during division. Here we show that a polarized membrane reservoir constructed by the NSC in early mitosis provides the source for expansion. The reservoir is formed from membrane domains that contain folds and microvilli that become polarized by apically-directed cortical flows of actomyosin early in mitosis. When furrow ingression begins and internal pressure increases, the stores of membrane within the apical reservoir are rapidly consumed. Expansion is significantly diminished in NSCs that lack a reservoir, and membrane expansion equalizes when the reservoir is not polarized. Our results suggest that the cortical flows that remodel the plasma membrane during asymmetric cell division function to satisfy the dynamic surface area requirements of unequally dividing cells.

## Introduction

Cellular resources can be partitioned unequally during division to facilitate differences in sibling cell fate and proliferation potential. For example, dividing *Drosophila* neural stem cells (NSCs) direct most cell volume into the sibling that retains the NSC fate and remains highly proliferative. In contrast, the neural precursor (NP) sibling that is destined towards differentiation and a postmitotic state receives a small fraction of the total volume (Gallaud et al., 2017; Homem and Knoblich, 2012). While asymmetric spindle positioning can contribute to unequal cell division, the size asymmetry of the NSC and NP cells results from preferential expansion of the nascent NSC surface during anaphase (Cabernard et al., 2010; Connell et al., 2011; Pham et al., 2019; Roubinet et al., 2017). Biased expansion is characterized by a rapid increase in surface area near the NSC pole while the area near the other pole remains relatively constant (Connell et al., 2011). Expansion occurs when intracellular hydrostatic pressure increases following the initiation of furrow ingression (Pham et al., 2019). While a biased increase in surface area is a key aspect of the unequal NSC division, little is known about how the plasma membrane accommodates asymmetric polar expansion. Here we examine the plasma membrane dynamics that support the geometric changes that accompany the unequal NSC division.

Morphological changes can significantly alter cell surface area, placing significant demands on the plasma membrane. Many animal cells, including the NSC, become spherical before dividing, a geometry that uses surface area most efficiently for a given volume. Division can lead to rapid morphology changes, including elongation and furrow ingression, that can quickly increase surface area as the cell assumes a non-spherical shape. However, the plasma membrane is relatively inelastic and can only undergo limited stretching to accommodate increases in surface area. One mechanism that cells use to buffer changes in surface area is to store membrane in cell surface reservoirs (Figard et al., 2013; Figard and Sokac, 2014; Kabeche et al., 2015). Membrane reservoirs use folds and microvilli to store membrane that is unnecessary for encapsulating the cell (i.e. excess membrane). Changes in cell shape that increase surface area, like those that accompany division, can be accommodated by tapping into the reservoir, in turn reducing the amount of excess membrane.

The NSC undergoes large, highly asymmetric morphological changes during division. Although it is not known to contain a reservoir, the NSC membrane is very dynamic, undergoing a polarity cycle that resembles its well-characterized protein polarity cycle (LaFoya and Prehoda, 2021). During the protein cycle, polarity proteins and fate determinants become polarized to the NSC’s apical and basal hemispheres during early mitosis (Gallaud et al., 2017; Homem and Knoblich, 2012; Knoblich, 2010, 2008). The apical proteins are polarized in part by apically-directed cortical flows of actomyosin. These cortical movements lead to the formation of an organized cap near the apical pole that remains until it depolarizes shortly before division (Oon and Prehoda, 2021, 2019). The plasma membrane undergoes a similar cycle in which membrane features that are initially dispersed over the surface of the cell move apically, in synchrony with polarity proteins, before appearing to spread over the surface by moving basally during anaphase (LaFoya and Prehoda, 2021). The dynamic NSC membrane features can be detected with diverse membrane markers and some are clearly microvilli suggesting that they could store excess membrane. Here we examine whether the NSC membrane contains a reservoir and if so, how membrane dynamics might contribute to the biased expansion that mediates unequal NSC division.

## Results

### The apparent surface area of the apical NSC sibling increases during division

We first estimated the demand on the plasma membrane during the divisions of *Drosophila* larval brain NSCs (Figure 1A). Like many animal cells, NSCs become spherical in early mitosis and the sibling cells remain spherical following the completion of furrowing (Figure 1B). The difference between the surface area required to encapsulate the NSC before division (e.g. at metaphase) and that for both siblings represents the amount of additional plasma membrane surface area required for the division. We term the surface area required to encapsulate the cell the *apparent surface area*, as the plasma membrane can store a reservoir of additional surface area in membrane fine structure such as folds and microvilli, which we term *excess membrane* (Figure 1C). We measured the overall increase in apparent surface area using Miranda-GFP, which marks the surface area of the nascent NP and NP siblings directly (with membrane-bound signal), and the nascent NSC and NSC sibling indirectly (with cytoplasmic signal; Figure 1D) (Ramat et al., 2017). We found that the apparent surface area increased by an average of 74 ± 19 μm^2^ (n = 10; mean ± SD) from metaphase to the completion of furrowing. We also used this method to estimate the bias in apparent surface area change between NSC and NP siblings, with the caveat that Miranda-GFP’s demarcation between NSC and NP may change slightly over this time frame. Consistent with previous measurements using the change in cell length along the division axis (Connell et al., 2011; Pham et al., 2019; Roubinet et al., 2017), we found that expansion is highly asymmetric, with an average increase of 94 ± 17 μm^2^ for the NSC apparent surface area and a relatively small but significant decrease of -20 ± 10 μm^2^ for the NP cell (n = 10; Figure 1E). The decrease may arise from shrinking of the Miranda-GFP domain but nevertheless indicates that the overall increase in surface area is mediated primarily by expansion of the nascent NSC surface. We conclude that the NSC division creates a significant demand for new surface area that is satisfied by a biased expansion process that directs the increase in surface area primarily towards the future NSC sibling.

**Figure 1.**
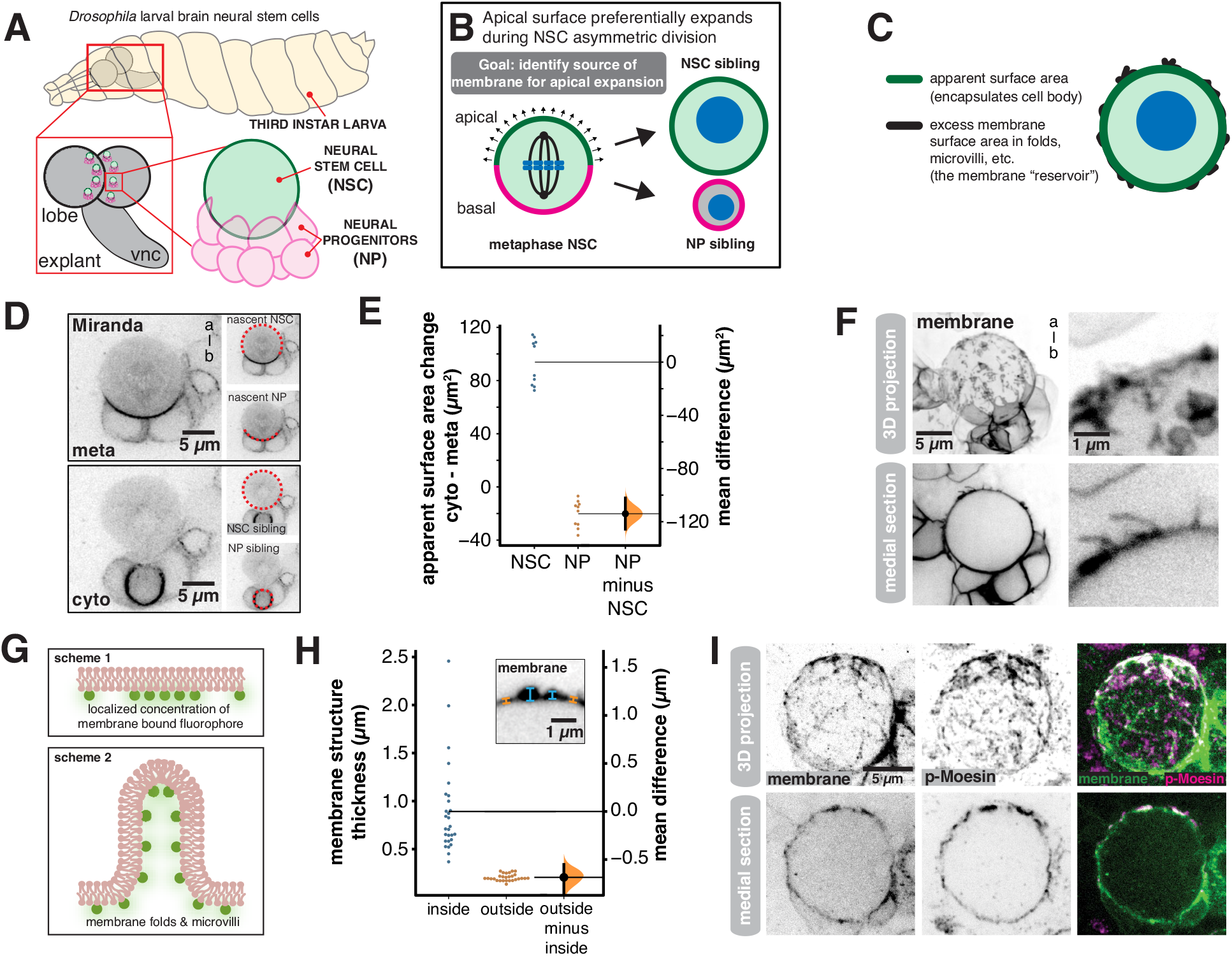
Neural stem cell membrane domains store excess membrane (A) *Drosophila* larval central nervous system explants used in this study. Larval brain lobe neural stem cells were imaged from explants dissected from third instar larvae (vnc: ventral nerve cord). (B) Model for the unequal division of neural stem cells. Preferential expansion of the apical hemisphere during anaphase contributes to size asymmetry. The goal of this study was to identify the membrane source for biased expansion. NSC = neural stem cell. NP = neural progenitor. (C) Apparent surface area and excess membrane. The total plasma membrane surface area is the sum of the surface area required to encapsulate the cell body (“apparent surface area”) and the surface area of membrane structures such as folds and microvilli (“excess membrane”). (D) Use of the NP marker Miranda (Miranda-GFP expressed from its endogenous locus) to measure NSC and NP apparent surface areas from metaphase (“meta”; immediately prior to cell elongation) and the completion of observable furrowing (“cyto”; cytokinesis). Dashed red lines show the demarcation of NSC and NP and “a-b” denotes apical-basal axis. A medial section of the NSC along the apical-basal axis is shown for each time point. (E) Change in apparent surface area that accompanies NSC division. The difference in apparent surface area from metaphase (“meta”) to the completion of observable furrowing (“cyto”; cytokinesis) is shown for divisions of 10 different NSCs using Miranda-GFP to mark the NSC and NP sibling surfaces. (F) Examples of NSC membrane structure associated with localized regions of increased membrane marker intensity (“a-b” denotes apical-basal axis). The membrane marker is expressed in neural stem cells and their progeny (worniu-GAL4>UAS-farnesyl-GFP). A three- dimensional projection of the front hemisphere of the NISC and adjacent progeny cells is shown (top-left panel) adjacent to a close up of the NSC surface (top-right panel). A medial section along the apical-basal axis is shown in the bottom-left adjacent to a close up of the NSC surface (bottom-right). (G) Schemes for localized enrichment of membrane marker: 1) a configuration that yields a localized increase in membrane marker concentration by increased density on a single plasma membrane. 2) a configuration that yields a localized increase in intensity with uniform membrane marker concentration but increased membrane density. (H) Variation of NSC membrane thickness associated with localized regions of increased membrane marker intensity (worniu-GAL4>UAS-farnesyl-GFP; “membrane”). The structural thickness of the membrane at regions inside and outside of membrane domains (n = 28) is plotted in a Gardner-Altman estimation plot. The 95% confidence interval is shown along with the bootstrap estimation distribution. The inset depicts an example structural thickness measurement. (I) Colocalization of activated Moesin with membrane structures. A neural stem cell expressing the membrane marker worniu-GAL4>UAS-PLCδ-PH-GFP fixed and stained with anti-GFP (“membrane”) and anti-activated Moesin (“p-Moesin”; phosphorylated Moesin). A three- dimensional projection of the front hemisphere of the NSC is shown.

### Neural stem cell membrane domains are a reservoir of excess membrane

We sought to understand how the NSC satisfies the requirement for additional surface area during division and the mechanism by which it is biased towards the NSC rather than the NP sibling. The NSC membrane is heterogeneous, with distinct domains characterized by localized areas of relatively high membrane marker intensity (Figure 1F) (LaFoya and Prehoda, 2021). NSC membrane domains are distributed across the cell surface in interphase and early mitosis and can be detected with diverse membrane markers. The higher marker intensity within NSC membrane domains could arise from an increased density of specific membrane components in a single bilayer (Figure 1G-scheme 1). Alternately, increased amounts of plasma membrane in the form of folds or microvilli could also lead to higher marker intensity (Figure 1G-scheme 2), and some microvilli are known to be present (LaFoya and Prehoda, 2021). We examined the prevalence of excess membrane in membrane domains by measuring their thickness using super resolution imaging (Azuma and Kei, 2015). NSC membrane domains in regions without signal from surrounding cells were nearly exclusively associated with increased thickness relative to adjacent membrane, which had a relatively uniform thickness approximately equal to the diffraction limited width (Figure 1H). Furthermore, many membrane domains colocalized with activated Moesin, a marker for microvilli (Fehon et al., 2010) (Figure 1I). We conclude that excess membrane is stored on the NSC surface in discrete regions that are thicker than surrounding regions and often contain microvilli. The presence of excess membrane makes the total NSC membrane surface area greater than the apparent surface area of the cell body, a phenomenon that has been compared to the “coastline paradox” in which the size of an interface is influenced by the presence of structure at different length scales (Sokac, 2017).

### Excess plasma membrane moves apically to form a polarized membrane reservoir

NSC membrane domains that are dispersed over the cell surface in early mitosis become concentrated around the apical pole by metaphase (Figure 2A and Video S1) (LaFoya and Prehoda, 2021). Given our finding that membrane domains store excess membrane, we hypothesized that the polarization process causes excess membrane to accumulate specifically to the apical pole. Consistent with this hypothesis, we measured a significant increase in apical membrane structure thickness at metaphase compared to prophase (Figure 2B). We did not detect any basally-directed movement of excess membrane during this time period, nor did we detect an increase in the membrane structure’s thickness at the basal pole (Figure 2B, Video S1). We also observed deformation of domains along the apical-basal axis as they moved towards the apical pole (c.f. Figure 2A inset, Video S1), suggesting the presence of tension-induced deformation of the membrane structures. Cell contacts between the NSC and adjacent progeny cells also undergo a similar pattern of deformation along the apical-basal axis indicating the presence of extensive mechanical forces which polarize the NSC surface in early mitosis (Figure 2A and Video S1) (LaFoya and Prehoda, 2021). We conclude that the polarization of the NSC membrane occurs at least in part from accumulation of excess membrane near the apical pole due to directional transport from around the cell surface.

**Figure 2.**
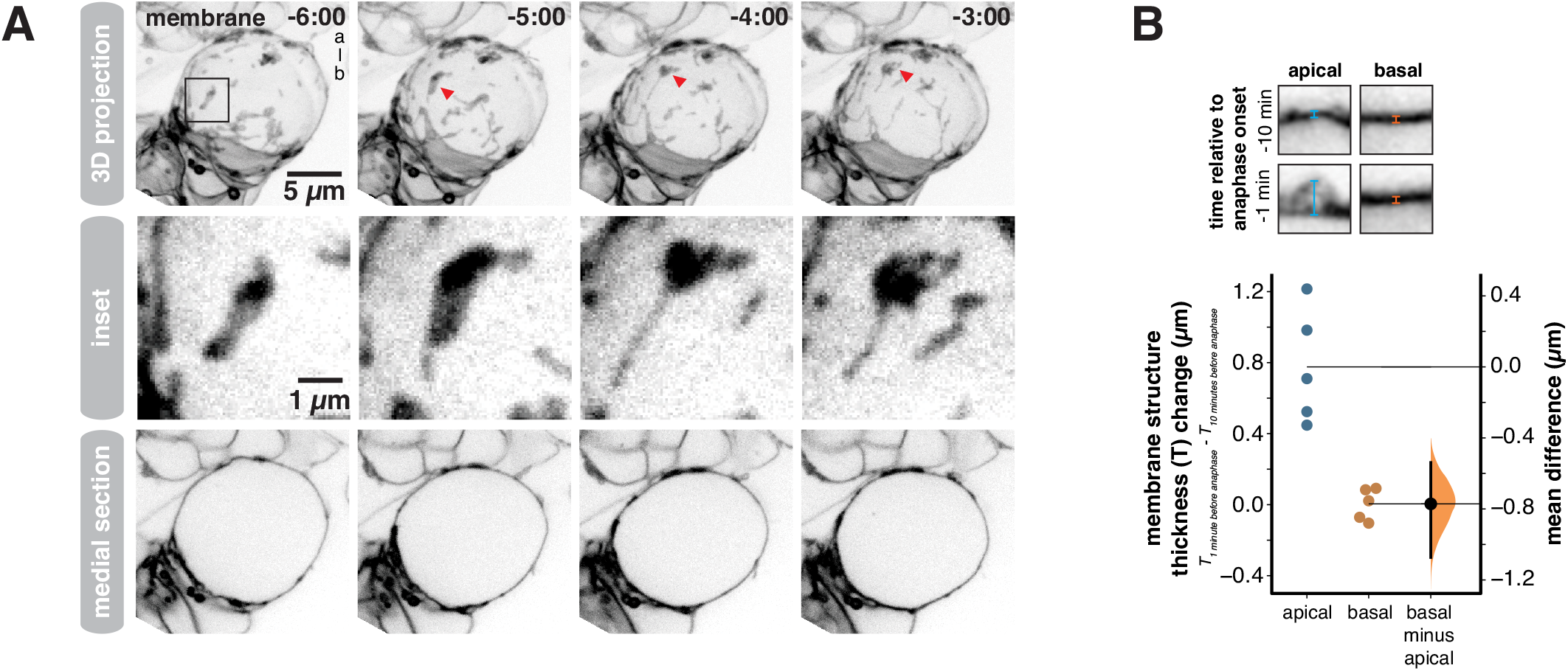
Apically directed membrane domain movements during prophase form an apical membrane reservoir (A) Frames from Video S1 showing the apically directed movement of excess membrane during prophase that leads to the accumulation of excess membrane near the apical pole at metaphase. Membrane is marked by worniu-GAL4>UAS-PLCδ-PH-GFP. The first row shows a three-dimensional projection for each time point, with arrow heads tracking one membrane domain as it moves apically. The second row shows an inset (box in first row) highlighting the membrane domain tracked in the first row. The third row shows a medial section for each time point. Time is shown relative to anaphase onset (m:ss) and “a-b” denotes the apical-basal axis. (B) Change in membrane structure thickness increases at the apical and basal poles between ten minutes and one minute prior to anaphase onset. A Gardner-Altman estimation plot is shown for the change in membrane thickness structure for apical and basal poles (n = 5 NSCs). The 95% confidence interval is shown along with the bootstrap estimation distribution. Insets show example measurements of membrane thickness at each pole.

### Cortical actomyosin flows drive apical plasma membrane accumulation

The apically-directed membrane domain movements that construct the polarized membrane reservoir do not occur in Latrunculin A-treated NSCs (LaFoya and Prehoda, 2021), indicating that they require F-Actin. Furthermore, apically-directed cortical flows of actomyosin in prophase NSCs have recently been reported (Oon and Prehoda, 2021), suggesting that cortical flows could drive the formation of the apical membrane reservoir. We examined whether cortical cytoskeletal flows are correlated with membrane polarization dynamics using super resolution, simultaneous imaging of the membrane and F-Actin, and separately membrane and Myosin II (Figure 3). As previously reported (LaFoya and Prehoda, 2021), we observed enrichment of F-Actin at sites of excess membrane (Figure 3 and Video S2). Here we find that the movements that polarize membrane domains are highly correlated with those of the cortical F-Actin meshwork that lies outside of the domains (Figure 3A,B and Video S2). We also examined the dynamics of Myosin II and did not observe its colocalization with membrane domains (Figure 3C and Video S3). However, the apically-directed movements of cortical Myosin II during prophase were strongly correlated with membrane domain movements that form the apical reservoir (Figure 3C,D and Video S3).

**Figure 3.**
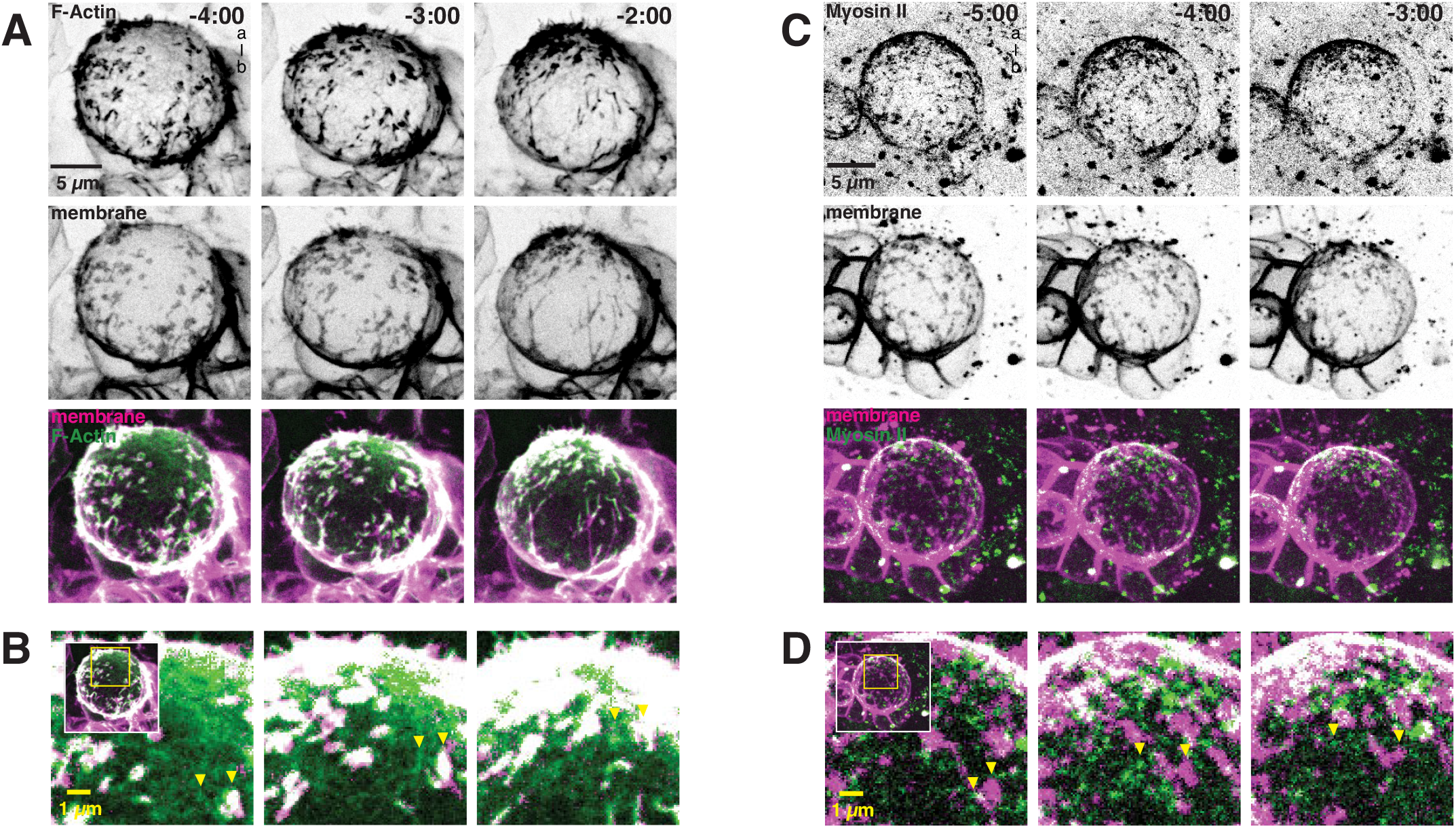
Apically directed flows of cortical actomyosin are correlated with the movement of excess membrane towards the apical pole (A) Selected frames from Video S2 showing coordinated movement of cortical F-Actin (worniu- GAL4>UAS-GMA [GFP-Moesin Actin binding domain]; “F-Actin”) and excess membrane domains (worniu-GAL4>UAS-PLCδ-PH-mCherry; “membrane”). A three-dimensional projection is shown for each time point. Time is shown relative to anaphase onset (m:ss) and “a-b” denotes apical-basal axis. (B) Coordinated movements of cortical Actin and membrane domains. Inset as shown from panel A with arrows highlighting moving membrane domain and cortical Actin mesh. (C) Selected frames from Video S3 showing coordinated movement of Myosin II (Sqh-GFP [Spaghetti Squash; *Drosophila* Myosin II regulatory light chain]; “Myosin II”) and excess membrane domains (worniu-GAL4>UAS-PLCδ-PH-mCherry; “membrane”). A three- dimensional projection is shown for each time point. Time is shown relative to anaphase onset (m:ss) and “a-b” denotes apical-basal axis. (D) Coordinated movements of cortical Myosin II and membrane domains. Inset as shown from panel C with arrows highlighting moving membrane domain and cortical Myosin II puncta.

The correlated motions of actomyosin and membrane suggest that actomyosin cortical flows are responsible for constructing the apical membrane reservoir. We tested this model by inhibiting Myosin II with the Rho Kinase inhibitor Y27632. While Y27632 can also inhibit the polarity kinase aPKC (Atwood and Prehoda, 2009), aPKC activity is not required for apical movements of membrane domains (LaFoya and Prehoda, 2021). We found that addition of Y27632 inhibited domain movement leading to failure to establish the apical membrane reservoir (Figure 4A-C and Video S4). Our results indicate that NSC membrane domains contain Actin and activated Moesin (Figure 1I), but not Myosin II (Figure 3C), and that the domains move apically during prophase with dynamics that are highly correlated with actomyosin cortical flows and that require Myosin II activity (Figure 4D).

**Figure 4.**
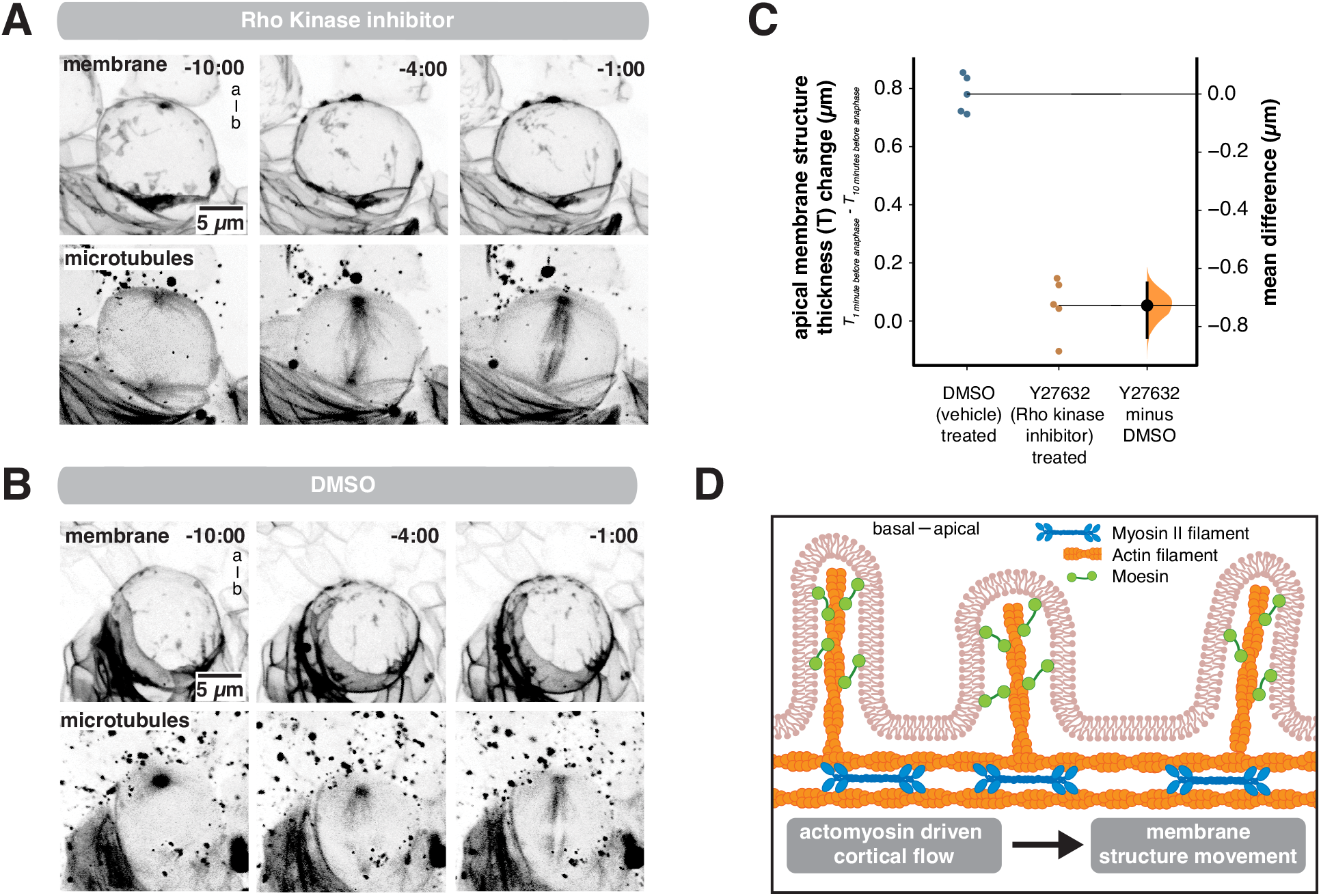
Activity of the Myosin II regulator Rho Kinase is required for NSC membrane polarization (A) Frames from Video S4 showing the effect of the Rho Kinase inhibitor (Y27632) on membrane reservoir formation. Membrane is marked by worniu-GAL4>UAS-PLCδ-PH-GFP and microtubules are marked by worniu-GAL4>UAS-Zeus-mCherry. A three-dimensional projection is shown for each time point. Time is shown relative to anaphase onset (mm:ss) and “a-b” denotes apical-basal axis. (B) Frames from Video S4 showing the effect of the vehicle DMSO on membrane reservoir formation. Membrane is marked by worniu-GAL4>UAS-PLCδ-PH-GFP and microtubules are marked by worniu-GAL4>UAS-Zeus-mCherry. Time is shown relative to anaphase onset (m:ss) and “a-b” denotes apical-basal axis. (C) Rho kinase activity is required for the increase in apical membrane thickness. A Gardner- Altman estimation plot is shown for the change in apical membrane thickness in treated (vehicle + Y27632) and untreated (vehicle only) cells. The 95% confidence interval is shown along with the bootstrap estimation distribution. (D) Model for membrane reservoir polarization. Excess membrane is contained in membrane fine structure such as microvilli that contain F-Actin and activated Moesin. Apically-directed flows of the cortical actomyosin cytoskeleton lead to coordinated movement of excess membrane domains towards the apical pole.

### Consumption of apical membrane reservoir during NSC division

The NSC constructs a polarized membrane reservoir in early mitosis by moving excess membrane to the apical pole using actomyosin driven cortical flows. How might the NSC use this structure to accommodate the rapid morphological changes that occur in late mitosis? We examined the dynamics of the apical membrane reservoir during anaphase, when the apical region’s apparent surface area increases (Figure 1E) (Connell et al., 2011; Pham et al., 2019). The internal pressure of the NSC changes dramatically during this phase of the cell cycle, peaking shortly after furrow ingression followed by a rapid decrease that coincides with membrane expansion (Pham et al., 2019). We followed the dynamics of the apical membrane reservoir during anaphase using super resolution imaging with high temporal resolution. We observed thinning of the apical membrane structure that coincided with the morphological changes associated with late mitosis (cell elongation and furrow ingression), and the increase in apical surface area (Figure 5 and Video S5; the full formation and consumption process is shown in Video S6). Microvilli that extended over a micron from the cell surface were reabsorbed into the cell as furrowing continued (Figure 5A and Video S5). As membrane reservoir consumption occurred, the apical surface expanded and the remaining domains of excess membrane appeared to move basally (Figure 5B), leading to a decrease in membrane structure thickness at the apical pole but no detectable change at the basal pole (Figure 5C). To ensure that the apical membrane structure thickness decrease was not simply a consequence of domains moving out of the focal plane, we monitored the maximum thickness of individual domains over time across optical sections and observed a significant decrease beginning at anaphase onset (Figure 5D). Both F-Actin (Figure 5E and Video S7) and Myosin II (Figure 5F and Video S8) migrated towards the cleavage furrow during this time period, as previously described (Roth et al., 2015; Roubinet et al., 2017). Thus, NSCs construct a polarized membrane reservoir in early metaphase that is consumed late in mitosis as the apical membrane expands.

**Figure 5.**
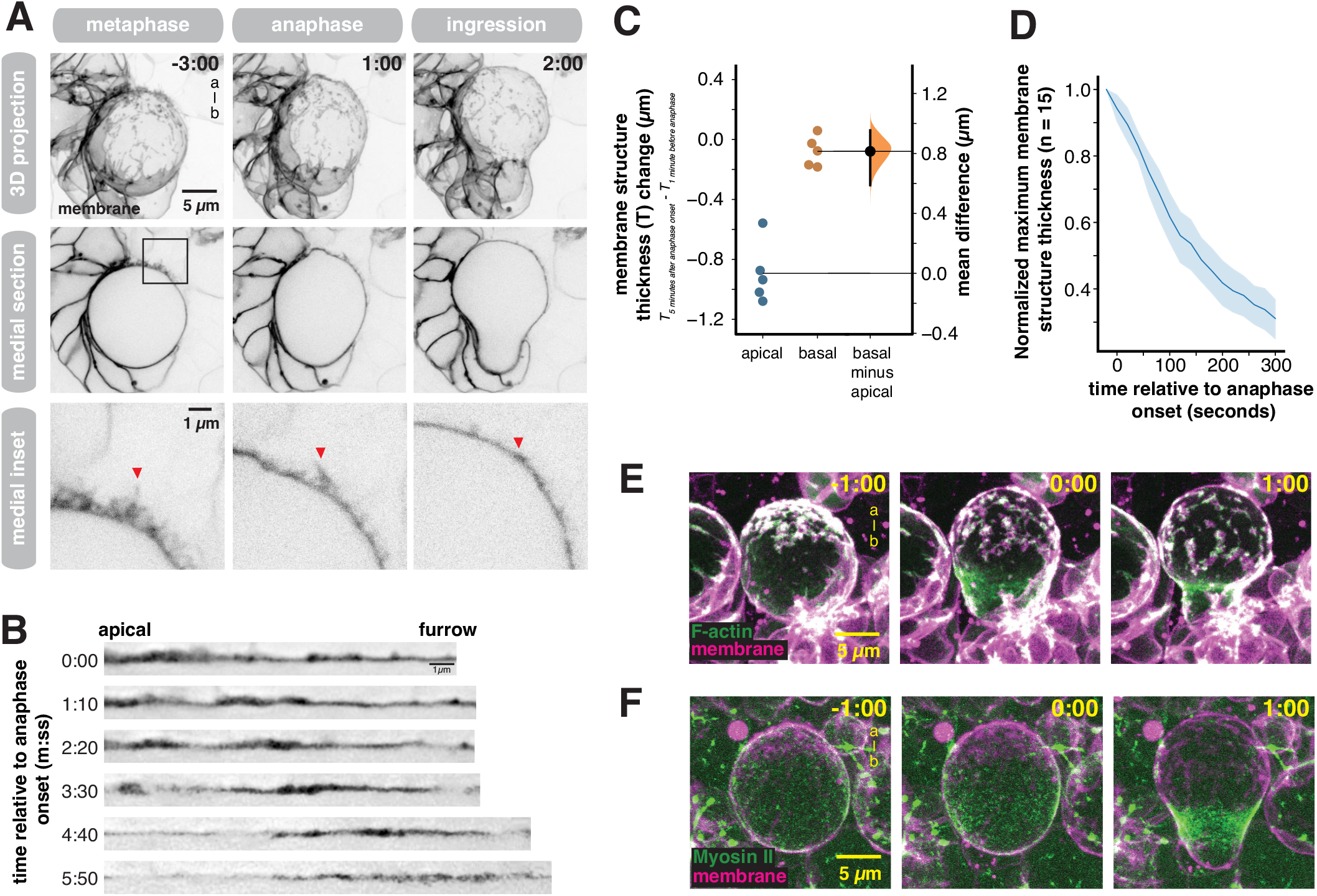
Consumption of the apical membrane reservoir during anaphase expansion (A) Frames from Video S5 showing the consumption of the apical membrane reservoir through anaphase and furrow ingression. Membrane is marked by worniu-GAL4>UAS-PLCδ-PH-GFP. The first row shows a three-dimensional projection, the second row shows a medial section for each time point, and the third row shows the inset specified by the box in the second row. Arrows show microvilli as it is absorbed into the cell body. Time is shown relative to anaphase onset (m:ss) and “a-b” denotes apical-basal axis. (B) Straightened membrane sections from Video S5 showing the apical pole to the furrow at times relative to anaphase onset. (C) Change in membrane structure thickness at the apical and basal poles between metaphase and the completion of furrowing. A Gardner-Altman estimation plot is shown for membrane structure thickness change at both locations. The 95% confidence interval is shown along with the bootstrap estimation distribution. (D) Membrane domain consumption dynamics during anaphase. The maximum membrane structure thickness (across x, y, and z) for individual domains (n = 15) is shown as a function of time relative to anaphase onset. The thickness is normalized to the starting value. The line represents the mean and the shaded region is one standard deviation. (E) F-Actin dynamics during anaphase membrane reservoir consumption. Selected frames from Video S7 showing coordinated movement of cortical F-Actin (worniu-GAL4>UAS-GMA; “F- Actin”) and excess membrane domains (worniu-GAL4>UAS-PLCδ-PH-GFP; “membrane”). Time is shown relative to anaphase onset (m:ss) and “a-b” denotes apical-basal axis. (F) Myosin II dynamics during anaphase membrane reservoir consumption. Selected frames from Video S8 showing coordinated movement of Myosin II (Sqh-GFP; “Myosin II”) and excess membrane domains (worniu-GAL4>UAS-PLCδ-PH-mCherry; “membrane”). Time is shown relative to anaphase onset (m:ss) and “a-b” denotes apical-basal axis.

### The membrane reservoir is required for expansion and completion of cytokinesis

Our data indicate that the apical membrane reservoir is consumed during anaphase, suggesting that the reservoir mediates preferential expansion at the apical pole. To investigate how the NSC division process proceeds without a reservoir, we examined the divisions of cyclodextrin-treated NSCs. Cyclodextrin treatment can ablate membrane domains in diverse cell types (Barman and Nayak, 2007; Pang et al., 2007), perhaps because it extracts cholesterol, significantly decreasing membrane rigidity (Chakraborty et al., 2020). In previous work, we found that cyclodextrin treatment inhibits formation of NSC membrane features (LaFoya and Prehoda, 2021). Consistent with previous observations, we found that the membranes of cyclodextrin-treated NSCs were relatively featureless such that the plasma membrane smoothly followed the contours of the cell (Figure 6A), apparently lacking any excess membrane. In the absence of a membrane reservoir, very little polar expansion occurred, and furrowing was incomplete and unstable (Figure 6B,C and Video S9). The failure to robustly furrow is consistent with a model in which the NSC reservoir provides the membrane necessary for the apparent surface area increase during division (Figard et al., 2016; Figard and Sokac, 2014). However, it is also possible that the cytokinesis defect arises at least in part from another effect of the cyclodextrin treatment, such as its effect on membrane rigidity.

**Figure 6.**
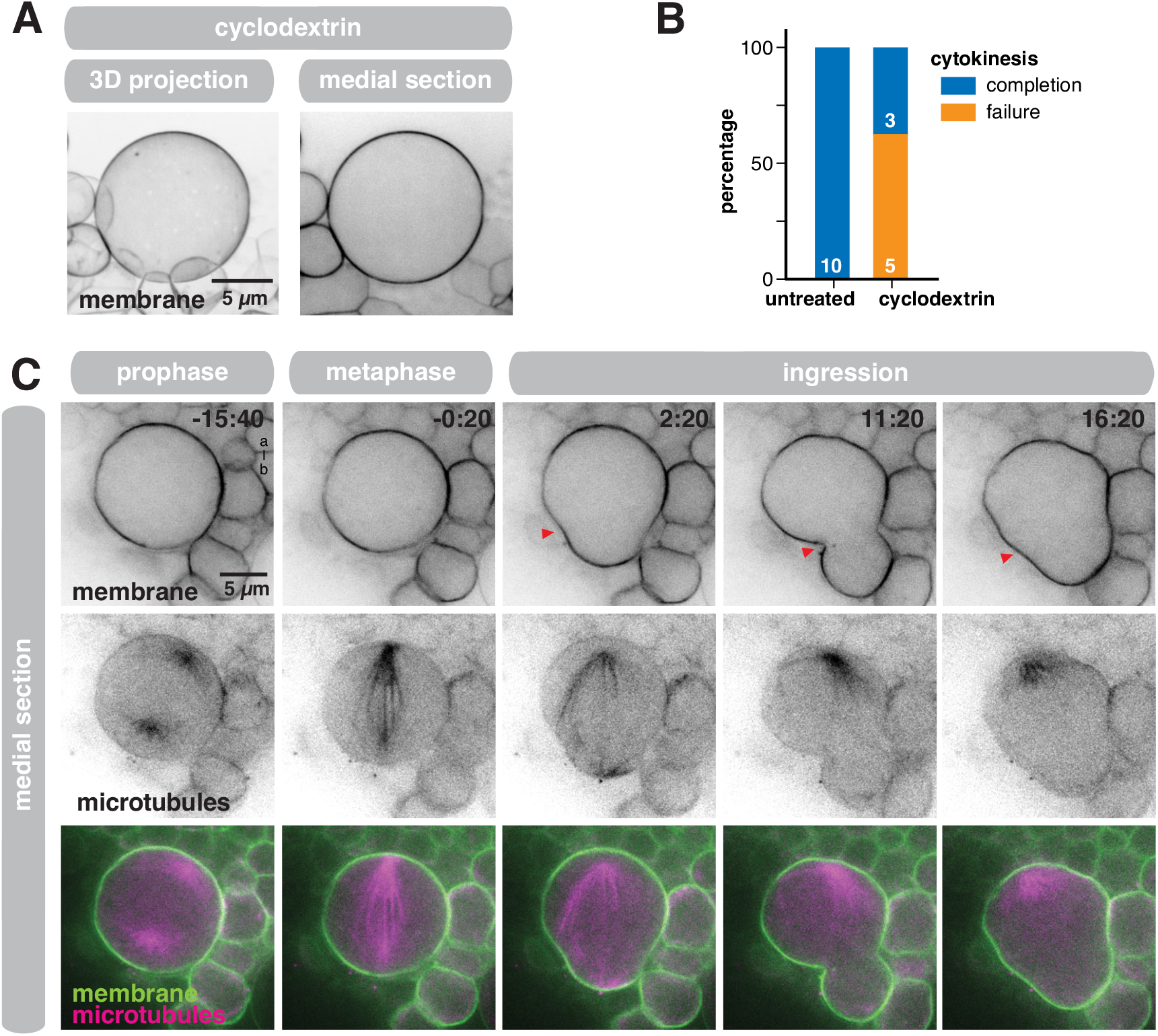
Cyclodextrin treatment removes excess membrane (A) The effect of Methyl-β-cyclodextrin (“cyclodextrin”) treatment on the NSC membrane (marked by worniu-GAL4>UAS-PLCδ-PH-GFP). A three-dimensional projection and a medial section is shown. (B) Effect of cyclodextrin treatment on cytokinesis. “Completion” denotes cells that completed furrowing whereas “failure” denotes cells whose furrow failed to completely ingress. Outcomes of cytokinesis for untreated NSCs are compared to cyclodextrin treated NSCs. (C) Selected frames from Video S9 showing the effect of cyclodextrin treatment on the NSC membrane and cytokinesis. Membrane is marked by worniu-GAL4>UAS-PLCδ-PH-GFP and microtubules are marked by worniu-GAL4>UAS-Zeus-mCherry. Arrows mark furrow that initially ingresses and ultimately retracts. Time is shown relative to anaphase onset (mm:ss) and “a-b” denotes apical-basal axis.

### Membrane reservoir polarization is required for biased expansion

Our data suggest that membrane reservoir polarization mediates biased expansion during NSC division. We examined how failure to polarize the membrane reservoir influences expansion using NSCs expressing RNAi directed against the heterotrimeric G-protein Gγ1 (hereafter Gγ). Gγ is known to be required for asymmetric NSC division (Izumi et al., 2004), and we found that, unlike wild-type NSCs, NSCs expressing Gγ RNAi do not polarize their membranes in early mitosis (Figure 7 and Video S10). In early mitosis, cells expressing Gγ RNAi had excess membrane distributed over their surface, like their wild-type counterparts. However, the apically-directed movements that polarize the membrane reservoir of wild-type NSCs failed to occur in Gγ RNAi-expressing cells. Without these polarizing movements, these cells failed to form a membrane reservoir at the apical pole as excess membrane remained distributed across the cell surface as furrow ingression commenced (Figure 7C). In the absence of a polarized membrane reservoir, the expansion that occurred following ingression was equally distributed to both poles and daughter cell size was approximately symmetric. We conclude that the biased expansion of the NSC membrane is mediated by polarization of the membrane reservoir before division.

**Figure 7.**
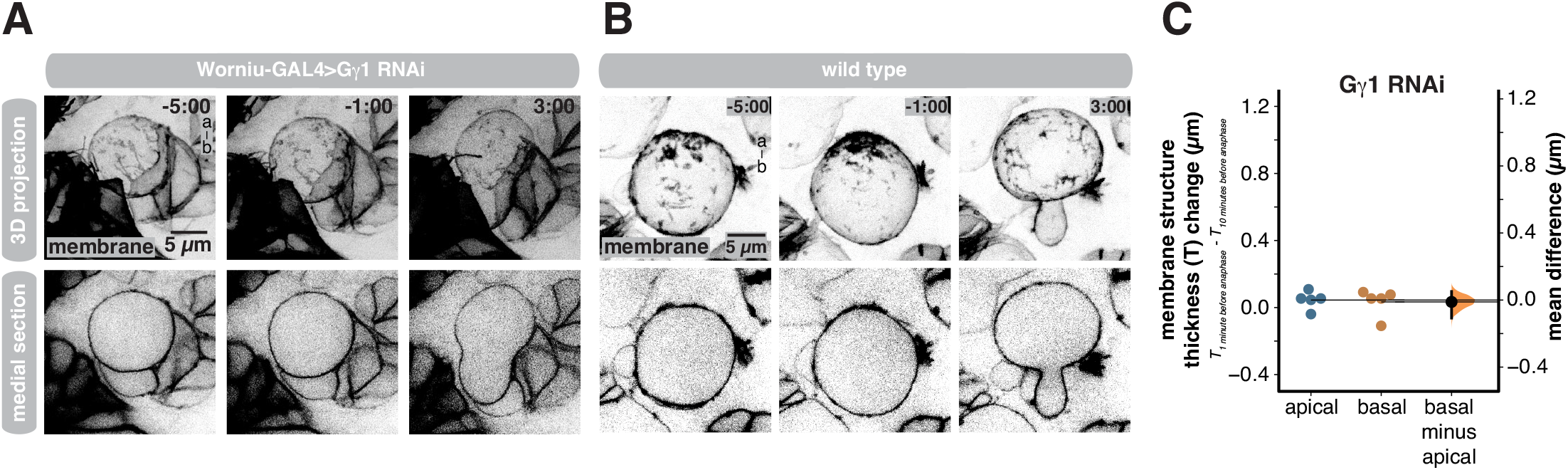
Membrane reservoir asymmetry is required for biased expansion (A) Selected frames from Video S10 of an NSC expressing RNAi directed against Gγ1 and the membrane marker UAS-PLCδ-PH-GFP (worniu-GAL4>UAS-PLCδ-PH-GFP; “membrane”). A three-dimensional projection and a medial section is shown for each time point. Time is shown relative to anaphase onset (m:ss) and “a-b” denotes apical-basal axis. (B) Selected frames from Video S6 of a wild type NSC expressing the membrane marker worniu-GAL4>UAS-PLCδ-PH-GFP for comparison. A three-dimensional projection and a medial section is shown for each time point. Time is shown relative to anaphase onset (m:ss) and “a-b” denotes apical-basal axis. (C) Membrane structure thickness changes at the apical and basal poles 10 minutes and 1 minute prior to anaphase onset in NSCs expressing RNAi directed against Gγi. A Gardner- Altman estimation plot is shown for the change in membrane structure thickness at the apical and basal poles (n = 5 NSCs). The 95% confidence interval is shown along with the bootstrap estimation distribution.

## Discussion

The morphological changes that accompany cell division can place significant demands on the plasma membrane by rapidly increasing total surface area. During the unequal divisions of *Drosophila* NSCs, the surface area increase is biased, with the nascent NSC surface expanding as the furrow ingresses while the NP surface remains relatively constant (Figure 1E) (Connell et al., 2011; Pham et al., 2019). In general, the demand for membrane can be buffered by reservoirs of excess membrane stored in folds and microvilli, or eisosomes in yeast (Figard et al., 2016; Figard and Sokac, 2014; Goudarzi et al., 2017; Kabeche et al., 2015). We discovered a membrane reservoir that mediates the biased expansion that occurs during NSC asymmetric division (Figure 8). The reservoir is constructed from localized regions of excess membrane that are dynamically collected at the apical pole by apically-directed cortical flows of actomyosin that begin and end in early mitosis (Figure 3) (Oon and Prehoda, 2021). When the apical membrane starts expanding as cell elongation and cleavage furrow ingression commence, the excess membrane in the reservoir begins incorporating into the membrane that encapsulates the cell body. Expansion and reservoir consumption coincide with the drop in internal hydrostatic pressure that occurs during anaphase (Pham et al., 2019). Our results indicate that NSC membrane structures play a specific role in accommodating the rapid surface area changes that occur in the late phases of mitosis, but it is important to note that they do not exclude a role for these structures in other processes, such as in mediating intercellular signaling like cytonemes (Kornberg, 2014).

**Figure 8.**
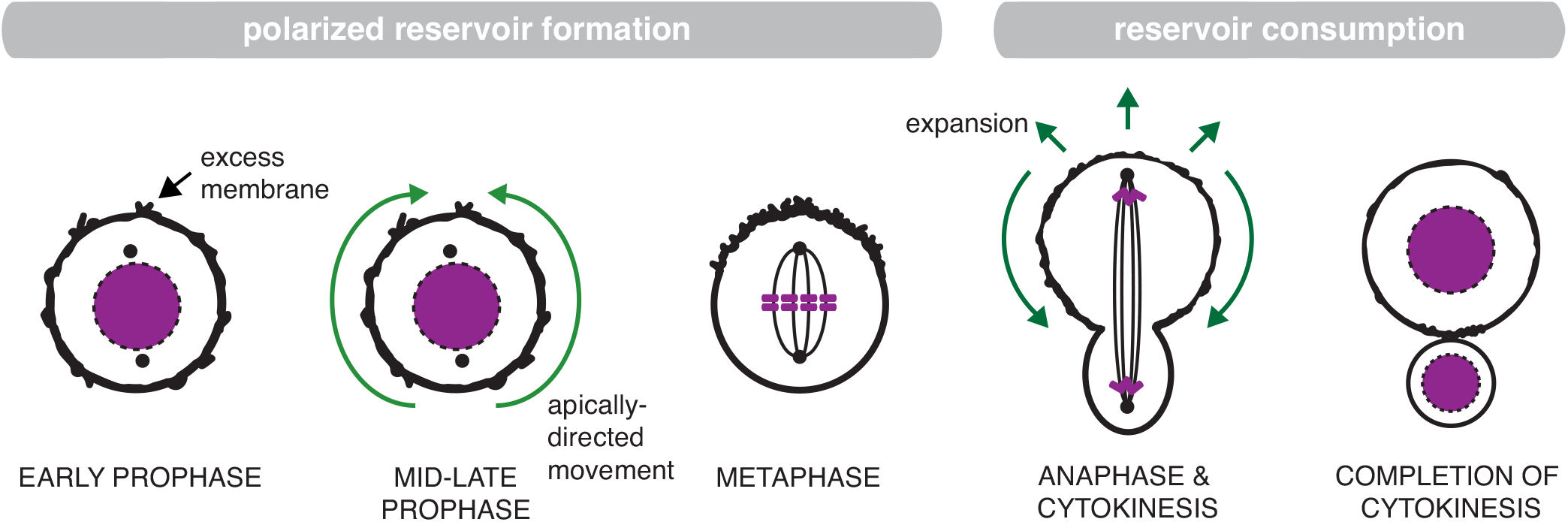
Model for reservoir-mediated asymmetric cell division In early mitosis, excess membrane contained within folds and microvilli are distributed across the cell surface. Apically-directed movements of membrane domains, mediated by directional flows of cortical actomyosin, lead to accumulation of the domains at the apical pole and the formation of an apical reservoir of excess membrane. The expansion that occurs at the apical pole, and creates the larger NSC sibling, is accompanied by consumption of the reservoir.

Examination of NSC membrane dynamics revealed that the cell goes to great lengths to prepare for biased expansion, generating directional cortical flows to collect excess membrane at the apical pole (Figure 2). Directional cortical flows of actomyosin play a role in the polarization of both the plasma membrane and proteins during the NSC division (LaFoya and Prehoda, 2021; Oon and Prehoda, 2021, 2019), and their role in cortical protein polarity of the asymmetrically-dividing *C. elegans* zygote has been extensively studied (Illukkumbura et al., 2019; Lang and Munro, 2017; Munro et al., 2004). Furthermore, polarized membrane structures have also been reported in the zygote (Hirani et al., 2019; Scholze et al., 2018), suggesting that they may play a more general role in asymmetric division. While cortical movements of polarity proteins such as aPKC are driven by actomyosin cortical flows in the NSC, F-Actin is not strictly required for its polarity. For example, in Latrunculin A treated NSCs that lack an Actin cytoskeleton, aPKC polarizes but ultimately diffuses into the basal hemisphere (Oon and Prehoda, 2019). Thus, currently available data suggest that actomyosin cortical flows in the NSC function primarily to organize the membrane and are not essential for establishing protein polarity, even while the cortical cytoskeleton maintains protein polarity by preventing spreading into the basal hemisphere (LaFoya and Prehoda, 2021; Oon and Prehoda, 2019).

How might actomyosin-generated cortical flows be coupled to the directional movement of excess membrane domains? Our results suggest a specific mechanism (Figure 4D). Membrane domains contain F-Actin that is coupled to the membrane via activated Moesin. This allows for direct coupling to the cortical cytoskeleton which undergoes apically-directed movements in early prophase (Figure 3) (Oon and Prehoda, 2021). This mechanism is similar to one described for cultured mammalian epithelial cells, in which microvilli move rapidly across the cell surface to form the brush border (Meenderink et al., 2019). Microvillar movement in both systems is based on coupling between F-Actin in microvilli and the cortical actomyosin cytoskeleton. In the developing brush border, however, microvillar motion is driven primarily by treadmilling of an interior Actin bundle. In contrast, movement of microvilli on the NSC surface is coupled to apically directed cortical flows of actomyosin (Figures 3,4). The NSC cortical flows are coupled not only to the movements of excess membrane, but also to the surrounding tissue (LaFoya and Prehoda, 2021; Oon and Prehoda, 2021), suggesting connections to a broad range of membrane-associated structures, including adhesions.

Shortly after its formation, the apical reservoir is tapped during anaphase (Figure 5), coinciding with the onset of furrow ingression and the peak of internal hydrostatic pressure (Pham et al., 2019). Simultaneously with these processes, Myosin II redistributes from the apical pole towards the cleavage furrow (Oon and Prehoda, 2021; Pham et al., 2019; Roubinet et al., 2017). Reservoir consumption could be caused by Myosin II relocalization or the initiation of furrow ingression, or a combination of both processes. Alternately redistribution of Myosin II could be a mechanical consequence of reservoir tapping.

## Acknowledgments

This work was supported by NIH grants GM127092 and F32GM134705.

## Star Methods

### Resource Availability

#### Lead Contact

Further information and requests for resources and reagents should be directed to the Lead Contact, Kenneth Prehoda.

#### Materials Availability

This study did not generate new unique reagents.

#### Data and Code Availability

The raw data supporting the current study have not been deposited in a public repository because of their large file size, but are available from the corresponding author on request.

### Experimental Model and Subject Details

#### Fly Strains

For tissue specific expression of UAS controlled transgenes in NSCs, a Worniu-GAL4 driver line was used. For live imaging of membrane dynamics, two different classes of membrane-bound fusion proteins were used, both of which nearly exclusively label the plasma membrane and not internal membranes in NSCs. UAS-farnesyl-GFP expresses the C-terminal region of human K-Ras tagged with GFP which becomes farnesylated and plasma membrane-anchored in cells. UAS-PLCδ-PH-GFP and UAS-PLCδ-PH-mCherry express the pleckstrin homology domain of human PLCδ tagged with GFP or mCherry, and binds to the plasma membrane lipid phosphoinositide PI(4,5)P_2_. F-Actin was imaged using UAS-GMA-GFP, which expresses the Actin binding domain of Moesin tagged with GFP. Progression through mitosis was tracked by expressing the microtubule marker UAS-Zeus-mCherry in NSCs, which labels microtubules allowing the visualization of the mitotic spindle. Symmetrical cell division in NSCs was induced through the expression of dsRNAi of G protein γ1 (Gγ1, FBgn0004921) under UAS control. Miranda-GFP expresses GFP tagged Miranda at the endogenous loci (Ramat et al., 2017).

Myosin II was imaged using GFP tagged Spaghetti squash (Sqh), the regulatory light chain of Myosin II, expressed from the natural promoter (Royou et al., 2002).

### Method Details

#### Live Imaging

Intact *Drosophila* central nervous systems were dissected from third instar larvae in a bath of Schneider’s Insect Media (SIM). These larval brain explants were then mounted dorsal side down on sterile poly-D-lysine coated 35mm glass bottom dishes (ibidi Cat#81156) containing modified minimal hemolymph-like solution (HL3.1). Explants were imaged on a Nikon Eclipse Ti-2 (60x H_2_O objective) equipped with a Yokogawa CSU-W1 SoRa spinning disk head and dual Photometrics Prime BSI sCMOS cameras. GFP tagged proteins were illuminated with 488nm. Proteins tagged with mCherry were illuminated with 561nm laser light. For super resolution imaging, individual NSCs were imaged using SoRa optics which achieve super resolution through optical photon reassignment. For time lapse super resolution imaging, images were acquired every 2-60 seconds. All figures and videos depict super resolution images, except in Video S10 where NSC #2 and #3 were captured using standard resolution imaging. For standard resolution time lapse imaging a z-stack of 41-61 optical sections with a step size of 0.5 μm was acquired every 10-20 seconds. For z-stacks acquired during super resolution imaging, a step size of 0.3 μm was used. Where time relative to anaphase onset is noted, anaphase onset was identified as the frame in which the apical membrane begins its expansion which coincides with the separation of the sister chromatids (LaFoya and Prehoda, 2021). To examine the effects of Rho Kinase inhibition on membrane dynamics, explants were treated with 1 mM Y27632 (solubilized in DMSO) for 5 minutes prior to live imaging. Explants treated with vehicle alone (DMSO only) were used for comparison. To examine the effects of cyclodextrin on membrane dynamics and cytokinesis, explants were treated with 15 mM methyl-β-cyclodextrin (solubilized in HL3.1) for 4 hours prior to live imaging. NSCs expressing PH-GFP and Zeus-mCherry and incubated with methyl-β-cyclodextrin (n=10) or left untreated (n=10) then imaged throughout mitosis and assessed by their ability to complete cytokinesis.

#### Immunofluorescent staining

To determine the localization of native activated Moesin in NSCs, the central nervous systems of third instar *Drosophila* larvae expressing Worniu-GAL4>UAS-PLCδ-PH-GFP were fixed in 4% paraformaldehyde. Mouse anti-GFP antibodies were used to stain neural stem cell membranes (marked by Worniu-GAL4> UAS-PLCδ-PH-GFP), and rabbit anti-activated Moesin antibodies were used to stain the phosphorylated form of Moesin. Primary antibodies were used at a concentration of 1:100. Alexa 488 labeled anti-mouse (Invitrogen) and Alexa 555 labeled anti-rabbit (Invitrogen) secondary antibodies were used to stain GFP labeled neural stem cell membranes and activated Moesin respectively. Super resolution images were captured using a Nikon Eclipse Ti-2 (60x H_2_O objective) equipped with a Yokogawa CSU-W1 SoRa spinning disk head and dual Photometrics Prime BSI sCMOS cameras.

#### Image Processing and Analysis

Images were analyzed using ImageJ (FIJI package) and Imaris (Bitplane) software. NSCs were identified through their location in the brain and large size, as well as the presence of transgenes expressed from the NSC specific promoter Worniu. 3D projections were created by z-stack processing in ImageJ, using the maximum intensity projection method for standard resolution imaging and standard deviation projection method for super resolution imaging. Only optical sections from the front hemisphere of the NSC were used to create 3D projections. We used a single optical section through the center of the cell along the apical-basal axis as medial sections. Photobleaching during time-lapse imaging was corrected for using ImageJ bleach correction tool in histogram matching mode. When image deconvolution was applied, deconvolution was performed using Nikon Elements standard 2D deconvolution mode. Image deconvolution was applied to Video S5 NSC #3. To reduce image noise, Gaussian blur was applied to the Zeus-mCherry channel in Figure 6C and Video S9 using ImageJ. Straightened membrane sections (Figure 5B) were prepared by manually tracing one side of the NSC membrane from the apical pole to the cleavage furrow and applying the straighten function in the ImageJ software package.

For the cell surface area quantifications in Figure 1E, cells expressing Miranda-GFP were used to calculate the surface area of metaphase NSCs and each sibling cell immediately following division. In ImageJ, a medial section was used to measure the diameter of the cell at five different angles from which an average radius was obtained to be used in the following calculations. The surface area was calculated with the equation SA = 4πr^2^, where SA = surface area and r = radius. From metaphase NSC images, the surface area of the nascent NP membrane was calculated using the equation 2πrh, where r = radius and h = height of the Miranda marked cap. The surface area of the nascent NSC membrane was calculated by subtracting the surface area of the nascent NP membrane from the total surface area of the metaphase NSC. The surface area of the nascent NSC at metaphase was compared to the surface area of the newly born NSC sibling immediately following division. Likewise, the surface area of the nascent NP at metaphase was compared to the surface area of the newly born NP sibling immediately following division.

To quantify the variations in thickness of the plasma membrane (Figure 1H), the thickness of various regions of the plasma membrane was measured in NSCs marked with two different membrane markers, UAS-farnesyl-GFP and UAS-PH-mCherry driven by Worniu-GAL4. Areas of the membrane which had high signal intensity in both channels (relative to other regions of the plasma membrane) were compared to areas of the membrane which exhibited average signal intensity. Medial sections from seven different NSCs were used to measure the thickness of the plasma membrane within areas of high signal intensity and also at four regions of average signal intensity for each cell (totaling n=28 for each). Gardner-Altman estimation plots for two group datasets and 95% confidence intervals of membrane thickness were prepared using the DABEST package (Ho et al., 2019).

To quantify the thickness of membrane structures over time (Figure 5D), a z-stack of neural stem cells expressing the membrane marker Worniu-GAL4>UAS-PLCδ-PH-GFP was acquired every 20 seconds. The thickness of individual membrane domains was measured every 20 seconds for 300 seconds total starting at anaphase onset. To account for movement of the membrane domain in the z-dimension, only the optical section with the thickest measurement at each time point was used for quantification. The fold change in membrane domain thickness for n = 15 membrane domains (n = 5 membrane domains from n = 3 neural stem cells from distinct brain explants) was averaged and plotted using matplotlib.

To quantify membrane thickness at the apical and basal poles (Figures 2B, 4C, 5C, and 7C) medial sections of NSCs were acquired using super resolution imaging. Images were loaded into ImageJ, and a line which traversed the cell membrane at either the apical or basal pole was drawn. If a cell junction between neighboring cells contacted the pole making it impossible to measure the thickness of the neural stem cell membrane, a line which traversed the cell membrane near to the pole but did not transect the cell junction was used. A plot profile was generated and the signal intensity peak correlating to the cell membrane was identified based on its position and high intensity over background signal. The length of the base of the peak was measured to determine the thickness of the cell membrane. Measurements were taken at various stages of the cell cycle and compared. When quantifying the effects of Gγ1 RNAi on membrane thickness the same protocol was used, but only NSCs which divided symmetrically were quantified. Gardner-Altman estimation plots for paired datasets and 95% confidence intervals of membrane thickness were prepared using the DABEST package (Ho et al., 2019) for n=5 NSCs from at least 3 separate brain explants.

#### Quantification and Statistical Analysis

Gardner-Altman estimation plots and 95% confidence intervals of datasets were prepared using the DABEST package (Ho et al., 2019). Statistical details can be found in the relevant methods section and figure legend.

## Key Resources Table

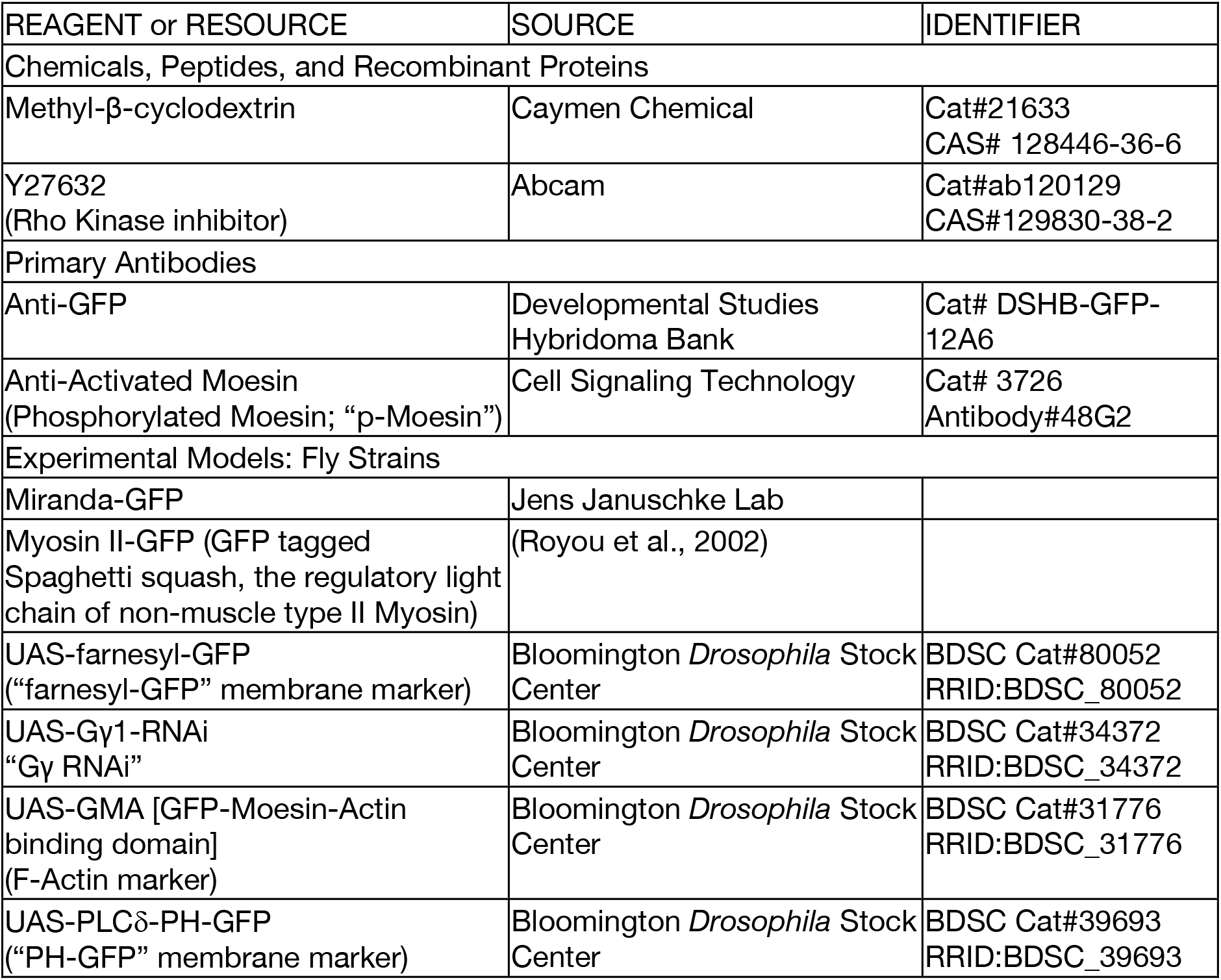
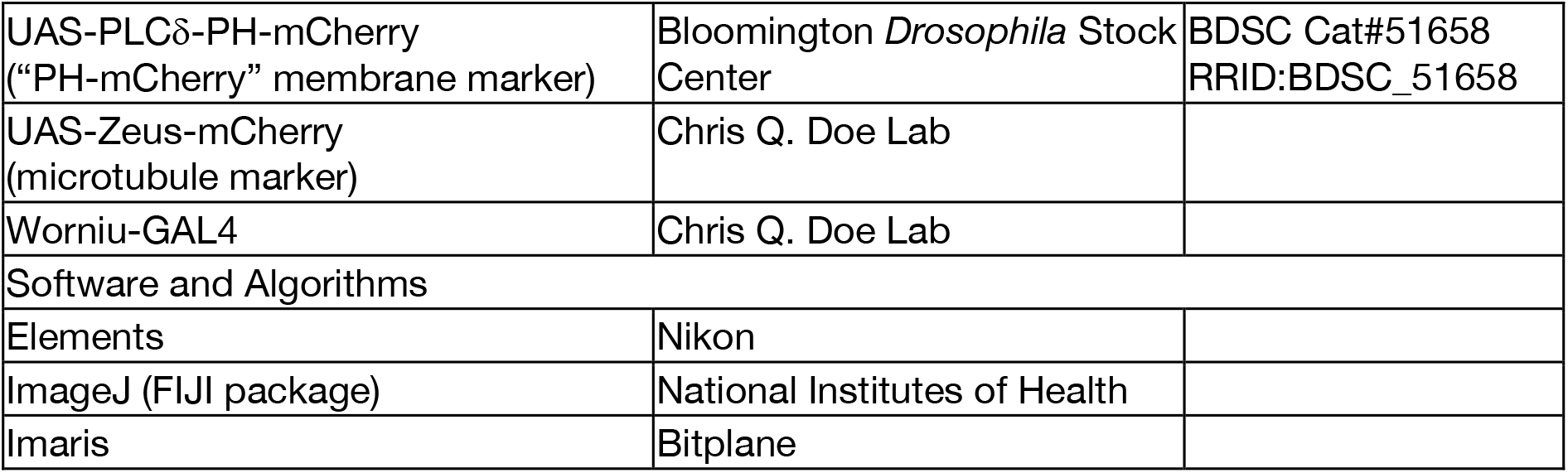

## Video Legends

Video S1: NSC membrane dynamics in early mitosis that lead to apical membrane reservoir formation

Super resolution videos of PH-GFP (worniu-GAL4>UAS-PLCδ-PH-GFP; “membrane”) expressing NSCs in early mitosis shown in 3D projections and in medial sections.

Video S2: Correlated NSC membrane and cortical Actin dynamics in early mitosis Super resolution videos of cortical F-Actin (worniu-GAL4>UAS-GMA; “F-Actin”) and PH-mCherry (worniu-GAL4>UAS-PLCδ-PH-mCherry; “membrane”) expressing NSCs in early mitosis shown in 3D projections and in medial sections.

Video S3: Correlated NSC membrane and Myosin II dynamics in early mitosis

Super resolution videos of GFP tagged Myosin type II (Myosin II) and PH-mCherry (worniu- GAL4>UAS-PLCδ-PH-mCherry; “membrane”) expressing NSCs in early mitosis shown in 3D projections and in medial sections.

Video S4: Membrane dynamics in Rho Kinase inhibitor treated neural stem cells

Super resolution videos of PH-GFP (worniu-GAL4>UAS-PLCδ-PH-GFP; “membrane”) and microtubules (worniu-GAL4>UAS-Zeus-mCherry; “microtubules”) expressing NSCs in early mitosis shown in 3D projections and in medial sections. Cells were treated with the Rho Kinase inhibitor, Y27632 (NSCs 1-3) or vehicle (DMSO) alone (NSCs 4-6).

Video S5: Consumption of the apical membrane reservoir during anaphase expansion Super resolution videos of PH-GFP (worniu-GAL4>UAS-PLCδ-PH-GFP; “membrane”) expressing NSCs during anaphase shown in 3D projections and in medial sections. “Close up views” of the apical membrane are provided in NSCs 1-3. Image deconvolution was applied to NSCs #3.

Video S6: NSC membrane dynamics during apical membrane reservoir formation and consumption

Super resolution videos of PH-GFP (worniu-GAL4>UAS-PLCδ-PH-GFP; “membrane”) expressing NSCs during the entirety of mitosis in 3D projections and in medial sections.

Video S7: NSC membrane and cortical Actin dynamics during apical membrane reservoir formation and consumption

Super resolution videos of cortical F-Actin (worniu-GAL4>UAS-GMA-GFP; “F-Actin”) and PH- mCherry (worniu-GAL4>UAS-PLCδ-PH-mCherry; “membrane”) expressing NSCs during the entirety of mitosis shown in 3D projections and in medial sections.

Video S8: NSC membrane and Myosin II dynamics during apical membrane reservoir formation and consumption

Super resolution videos of GFP tagged Myosin II and PH-mCherry (worniu-GAL4>UAS-PLCδ- PH-mCherry; “membrane”) expressing NSCs during the entirety of mitosis shown in 3D projections and in medial sections.

Video S9: Membrane dynamics in cyclodextrin-treated NSCs

Super resolution videos of PH-GFP (worniu-GAL4>UAS-PLCδ-PH-GFP; “membrane”) and microtubules (worniu-GAL4>UAS-Zeus-mCherry; “microtubules”) expressing NSCs during the entirety of mitosis shown in medial sections. Cells were treated with cyclodextrin (NSCs #1-4) or untreated (NSC #5).

Video S10: Membrane dynamics in Gγ RNAi expressing NSCs

Super resolution (NSC #1) and standard resolution (NSCs #2 & 3) videos of PH-GFP (worniu- GAL4>UAS-PLCδ-PH-GFP; “membrane”) and Gγ RNAi (worniu-GAL4>UAS-Gγ RNAi) expressing NSCs shown in 3D projections and in medial sections.

## References

Atwood SX, Prehoda KE. 2009. aPKC phosphorylates Miranda to polarize fate determinants during neuroblast asymmetric cell division. Curr Biol CB 19:723–729. doi:10.1016/j.cub.2009.03.056

Azuma T, Kei T. 2015. Super-resolution spinning-disk confocal microscopy using optical photon reassignment. Opt Express 23:15003–15011. doi:10.1364/OE.23.015003

Barman S, Nayak DP. 2007. Lipid raft disruption by cholesterol depletion enhances influenza A virus budding from MDCK cells. J Virol 81:12169–12178. doi:10.1128/JVI.00835-07

Cabernard C, Prehoda KE, Doe CQ. 2010. A spindle-independent cleavage furrow positioning pathway. Nature 467:91–94. doi:10.1038/nature09334

Chakraborty S, Doktorova M, Molugu TR, Heberle FA, Scott HL, Dzikovski B, Nagao M, Stingaciu L-R, Standaert RF, Barrera FN, Katsaras J, Khelashvili G, Brown MF, Ashkar R. 2020. How cholesterol stiffens unsaturated lipid membranes. Proc Natl Acad Sci U S A 117:21896–21905. doi:10.1073/pnas.2004807117

Connell M, Cabernard C, Ricketson D, Doe CQ, Prehoda KE. 2011. Asymmetric cortical extension shifts cleavage furrow position in Drosophila neuroblasts. Mol Biol Cell 22:4220–4226. doi:10.1091/mbc.E11-02-0173

Fehon RG, McClatchey AI, Bretscher A. 2010. Organizing the cell cortex: the role of ERM proteins. Nat Rev Mol Cell Biol 11:276–287. doi:10.1038/nrm2866

Figard L, Sokac AM. 2014. A membrane reservoir at the cell surface: unfolding the plasma membrane to fuel cell shape change. Bioarchitecture 4:39–46. doi:10.4161/bioa.29069

Figard L, Wang M, Zheng L, Golding I, Sokac AM. 2016. Membrane Supply and Demand Regulates F-Actin in a Cell Surface Reservoir. Dev Cell 37:267–278. doi:10.1016/j.devcel.2016.04.010

Figard L, Xu H, Garcia HG, Golding I, Sokac AM. 2013. The plasma membrane flattens out to fuel cell-surface growth during Drosophila cellularization. Dev Cell 27:648–655. doi:10.1016/j.devcel.2013.11.006

Gallaud E, Pham T, Cabernard C. 2017. Drosophila melanogaster Neuroblasts: A Model for Asymmetric Stem Cell Divisions. Results Probl Cell Differ 61:183–210. doi:10.1007/978-3-319-53150-2_8

Goudarzi M, Tarbashevich K, Mildner K, Begemann I, Garcia J, Paksa A, Reichman-Fried M, Mahabaleshwar H, Blaser H, Hartwig J, Zeuschner D, Galic M, Bagnat M, Betz T, Raz E. 2017. Bleb Expansion in Migrating Cells Depends on Supply of Membrane from Cell Surface Invaginations. Dev Cell 43:577-587.e5. doi:10.1016/j.devcel.2017.10.030

Hirani N, Illukkumbura R, Bland T, Mathonnet G, Suhner D, Reymann A-C, Goehring NW. 2019. Anterior-enriched filopodia create the appearance of asymmetric membrane microdomains in polarizing C. elegans zygotes. J Cell Sci 132. doi:10.1242/jcs.230714

Ho J, Tumkaya T, Aryal S, Choi H, Claridge-Chang A. 2019. Moving beyond P values: data analysis with estimation graphics. Nat Methods 16:565–566. doi:10.1038/s41592-019-0470-3

Homem CCF, Knoblich JA. 2012. Drosophila neuroblasts: a model for stem cell biology 139:4297–4310. doi:10.1242/dev.080515

Illukkumbura R, Bland T, Goehring NW. 2019. Patterning and polarization of cells by intracellular flows. Curr Opin Cell Biol 62:123–134. doi:10.1016/j.ceb.2019.10.005

Izumi Y, Ohta N, Itoh-Furuya A, Fuse N, Matsuzaki F. 2004. Differential functions of G protein and Baz-aPKC signaling pathways in Drosophila neuroblast asymmetric division. J Cell Biol 164:729–738. doi:10.1083/jcb.200309162

Kabeche R, Howard L, Moseley JB. 2015. Eisosomes provide membrane reservoirs for rapid expansion of the yeast plasma membrane. J Cell Sci 128:4057–4062. doi:10.1242/jcs.176867

Knoblich JA. 2010. Asymmetric cell division: recent developments and their implications for tumour biology. Nat Rev Mol Cell Biol 11:849–860. doi:10.1038/nrm3010

Knoblich JA. 2008. Mechanisms of asymmetric stem cell division. Cell 132:583–597. doi:10.1016/j.cell.2008.02.007

Kornberg TB. 2014. Cytonemes and the dispersion of morphogens. Wiley Interdiscip Rev Dev Biol 3:445–463. doi:10.1002/wdev.151

LaFoya B, Prehoda KE. 2021. Actin-dependent membrane polarization reveals the mechanical nature of the neuroblast polarity cycle. Cell Rep 35:109146. doi:10.1016/j.celrep.2021.109146

Lang CF, Munro E. 2017. The PAR proteins: from molecular circuits to dynamic self-stabilizing cell polarity. Dev Camb Engl 144:3405–3416. doi:10.1242/dev.139063

Meenderink LM, Gaeta IM, Postema MM, Cencer CS, Chinowsky CR, Krystofiak ES, Millis BA, Tyska MJ. 2019. Actin Dynamics Drive Microvillar Motility and Clustering during Brush Border Assembly. Dev Cell 50:545-556.e4. doi:10.1016/j.devcel.2019.07.008

Munro E, Nance J, Priess JR. 2004. Cortical flows powered by asymmetrical contraction transport PAR proteins to establish and maintain anterior-posterior polarity in the early C. elegans embryo. Dev Cell 7:413–424. doi:10.1016/j.devcel.2004.08.001

Oon CH, Prehoda KE. 2021. Phases of cortical actomyosin dynamics coupled to the neuroblast polarity cycle. eLife 10:e66574. doi:10.7554/eLife.66574

Oon CH, Prehoda KE. 2019. Asymmetric recruitment and actin-dependent cortical flows drive the neuroblast polarity cycle. eLife 8. doi:10.7554/eLife.45815

Pang DJ, Hayday AC, Bijlmakers M-J. 2007. CD8 Raft Localization Is Induced by Its Assembly into CD8αβ Heterodimers, Not CD8αα Homodimers. J Biol Chem 282:13884–13894. doi:10.1074/jbc.M701027200

Pham TT, Monnard A, Helenius J, Lund E, Lee N, Müller DJ, Cabernard C. 2019. Spatiotemporally Controlled Myosin Relocalization and Internal Pressure Generate Sibling Cell Size Asymmetry. iScience 13:9–19. doi:10.1016/j.isci.2019.02.002

Ramat A, Hannaford M, Januschke J. 2017. Maintenance of Miranda Localization in Drosophila Neuroblasts Involves Interaction with the Cognate mRNA. Curr Biol CB 27:2101-2111.e5. doi:10.1016/j.cub.2017.06.016

Roth M, Roubinet C, Iffländer N, Ferrand A, Cabernard C. 2015. Asymmetrically dividing Drosophila neuroblasts utilize two spatially and temporally independent cytokinesis pathways. Nat Commun 6:6551. doi:10.1038/ncomms7551

Roubinet C, Tsankova A, Pham TT, Monnard A, Caussinus E, Affolter M, Cabernard C. 2017. Spatio-temporally separated cortical flows and spindle geometry establish physical asymmetry in fly neural stem cells. Nat Commun 8:1383. doi:10.1038/s41467-017-01391-w

Royou A, Sullivan W, Karess R. 2002. Cortical recruitment of nonmuscle myosin II in early syncytial Drosophila embryos: its role in nuclear axial expansion and its regulation by Cdc2 activity. J Cell Biol 158:127–137. doi:10.1083/jcb.200203148

Scholze MJ, Barbieux KS, De Simone A, Boumasmoud M, Süess CCN, Wang R, Gönczy P. 2018. PI(4,5)P2 forms dynamic cortical structures and directs actin distribution as well as polarity in Caenorhabditis elegans embryos. Dev Camb Engl 145. doi:10.1242/dev.164988

Sokac AM. 2017. Seeing a Coastline Paradox in Membrane Reservoirs. Dev Cell 43:541–542. doi:10.1016/j.devcel.2017.11.013

